# The MAGOH paralogs - MAGOH, MAGOHB and their multiple isoforms

**DOI:** 10.1101/2021.05.07.443087

**Authors:** Ayushi Rehman, Pratap Chandra, Kusum Kumari Singh

## Abstract

A central processing event in eukaryotic gene expression is splicing. Concurrent with splicing, the core-EJC proteins, eIF4A3 and RBM8A-MAGOH heterodimer are deposited 24 bases upstream of newly formed exon-exon junctions. One of the core-EJC proteins, MAGOH contains a paralog MAGOHB, and this paralog pair is conserved across vertebrates. Upon analysis of the splice variants of MAGOH-paralogs, we have found the presence of alternate protein isoforms which are also evolutionarily conserved. Further, comparison of the amino acid sequence of the principal and alternate protein isoforms has revealed absence of key amino acid residues in the alternate isoforms. The conservation of principal and alternate isoforms correlates to the importance of MAGOH and MAGOHB across vertebrates.

## 1. INTRODUCTION

The primary step in eukaryotic gene expression involves transcription of DNA into precursor-mRNAs (pre-mRNAs). The pre-mRNAs undergo various post-transcriptional processing events before generating functional protein products. A notable processing event is splicing, where the spliceosome removes introns and ligates exons together. Concurrent with splicing, mature mRNAs are marked with the exon junction complex (EJC) at 24 nucleotides upstream of an exon-exon junction ^1^. The EJC forms a gene regulatory nexus composed of many proteins that are grouped under stable core- and transient peripheral-protein complexes. The core-EJC consists of three proteins, eIF4A3 and RBM8A-MAGOH heterodimer; whereas the peripheral-EJC includes ALYREF, ASAP complex or PSAP complex, and Upf proteins^2–6^. Additional to mRNA processing, the core-EJC proteins have essential functions in brain development and embryonic neurogenesis^7^. In this communication, we will mainly focus on one of the core-EJC proteins, MAGOH.

MAGOH (*mago nashi* homolog) was originally identified in *Drosophila melanogaster,* where embryos lacking functional mago protein produced sterile progeny. Thus, this protein was named “mago nashi” meaning “without grandchildren” in the Japanese language ^8^. MAGOH is well conserved among all vertebrates. It functions to maintain integrity of mRNAs as a part of the EJC as well as the nonsense-mediated mRNA decay (NMD) pathway^9^. The importance of MAGOH in NMD has recently also been characterised in zebrafish^10^. Interestingly, vertebrates contain a paralog of MAGOH, MAGOHB which is also associated with mRNAs via EJC and NMD^11^. In addition to their roles in mRNA processing, the MAGOH paralogs (MAGOH and MAGOHB) have been found to be associated with cancer progression. In cancers with a hemizygous deletion of MAGOH, MAGOHB was suggested as the highest dependent gene. The dependency of MAGOHB in such MAGOH deficient cancers have the potential for cancer treatment^12^. MAGOH paralogs have also been found to be involved in gastric cancers. Importantly, the simultaneous knockdown of MAGOH-MAGOHB in gastric cancer cells exhibited anti-tumor effects via the bRAF/MEK/ERK signalling pathway^13^. Thus, the MAGOH paralogs are essential for proper functioning of cells, yet sufficient information on the architecture and evolutionary conservation of these genes is lacking. Concordant with most eukaryotic genes, both the MAGOH-paralogs undergo alternative splicing and produce multiple splice variants, which have not yet been characterised. In this communication, we will discuss the features of the alternate protein isoforms of MAGOH-paralogs in humans and mice. We will also look at structural features of the alternate isoforms compared to the principal MAGOH protein isoform.

## 2. RESULTS and DISCUSSION

### 2.1 MAGOH paralogs have multiple protein isoforms in humans

MAGOH-paralogs belong to the MAGO NASHI protein family, Ensembl ID-PTHR12638. In humans, *MAGOH* also known as *MAGOH1* or *MAGOHA* is located on the reverse strand of chromosome 1 at 1p32.3 (*chr1:53,226,900-53,238,518, GRCh38/hg38*). Alternative splicing of *MAGOH* pre-mRNA generates four transcripts, *MAGOH-201, MAGOH-202, MAGOH-203*, and *MAGOH-204* (Table-1). *MAGOH-202* is the primary transcript made of five exons and encodes the principal protein isoform of 146 amino acids (aa). *MAGOH-201* constitutes four exons and codes for an alternate protein isoform of 109 aa. Comparison of *MAGOH-201* with *MAGOH-202* reveals skipping of exon 3 (111 nucleotides) in MAGOH-201, generating the alternate isoform (Figure-1a). The other transcripts, *MAGOH-203* and *MAGOH-204* comprise three exons and do not translate into proteins due to the absence of open reading frames (ORF). Hence, *MAGOH* is transcribed to generate four splice variants, of which two are protein-coding and two are non-coding. The finding of an alternate isoform (109 aa) is interesting as MAGOH is primarily mentioned as a 146 aa protein, with a molecular weight of 17 kDa ^14,15^. Subsequent to the finding of an alternate transcript in *MAGOH*, we were interested to find out if its paralog, *MAGOHB* also produces an alternate transcript.

**Figure 1-.**
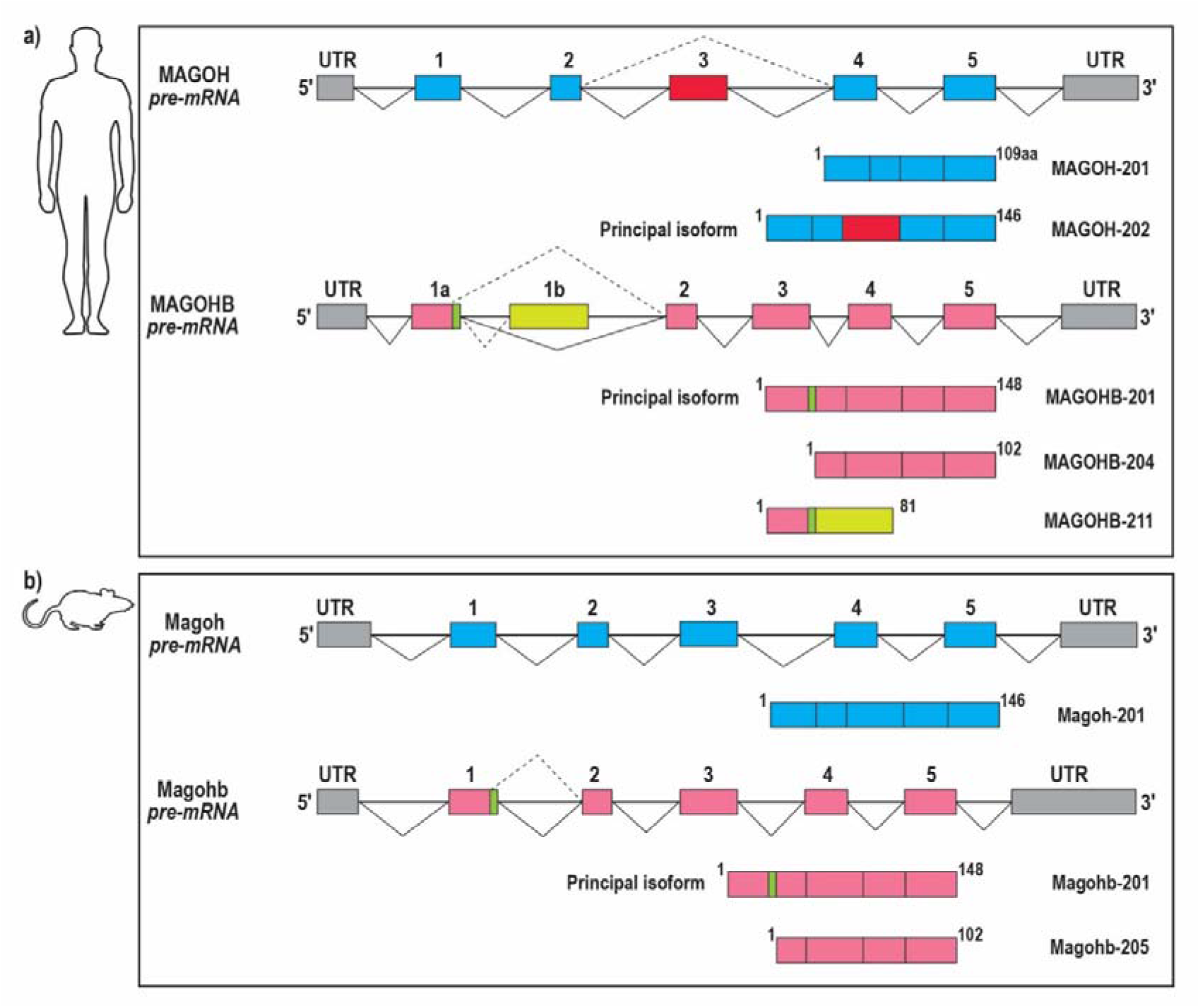
Isoforms of MAGOH-paralogs in (a) humans and (b) mice. The pre-mRNA generated from each gene is shown on top, where boxes represent exons, and introns are represented by horizontal lines. Constitutive splicing is denoted with solid lines, while alternative splicing is denoted with dashed lines. MAGOH is represented in blue and MAGOHB is represented in pink. The pre-mRNA shown here includes exons of the protein-coding transcripts. Principal and alternate protein isoforms are shown below each pre-mRNA, where each box represents coding exons from the pre-mRNA.

**Table 1-.** MAGOH paralog isoforms in *Homo sapiens*

| Transcript ID | Ensembl ID | Length (bases) | Protein (amino acids) | Biotype | 5'UTR (bases) | Exon 1 (bases) | Exon 2 (bases) | Exon3 (bases) | Exon4 (bases) | Exon5 (bases) | Exon6 (bases) | 3'UTR (bases) |
| --- | --- | --- | --- | --- | --- | --- | --- | --- | --- | --- | --- | --- |
| **MAGOH-201** | ENST00000371466.4 | 500 | 109 | Protein-coding | 51 | 51* + 88 | 59 | 83 | 100 + 119 | - | - | 119 |
| **MAGOH-202** | ENST00000371470.8 | 656 | 146 | Protein-coding | 70 | 70 + 88 | 59 | 111 | 83 | 100 + 145 | - | 145 |
| MAGOH-203 | ENST00000462941.1 | 738 | - | Processed transcript (no ORF) | - | 148 | 59 | 531 | - | - | - | - |
| MAGOH-204 | ENST00000495868.1 | 695 | - | Processed transcript | - | 398 | 83 | 214 | - | - | - | - |
| **MAGOHB-201** | ENST00000320756.7 | 2606 | 148 | Protein-coding | 77 | 77 + 94 | 59 | 111 | 83 | 100 + 2082 | - | 2082 |
| MAGOHB-202 | ENST00000398874.8 | 549 | - | IR* | - | 187 | 362 | - | - | - | - | - |
| MAGOHB-203 | ENST00000537852.5 | 2671 | - | IR | - | 162 | 59 | 2088 | - | - | - | - |
| **MAGOHB-204** | ENST00000539554.5 | 666 | 102 | Protein-coding | 95 | 51 | 44 + 15 | 111 | 83 | 100 + 262 |  | 262 |
| MAGOHB-205 | ENST00000540074.5 | 705 | 44 | NMD* | 95 | 51 | 45 + 15 | 111 | 9 + 277 | 83 | 115 | 475 |
| MAGOHB-206 | ENST00000543929.5 | 841 | 81 | NMD | 68 | 68 + 94 | 152 + 187 | 59 | 111 | 170 | - | 527 |
| MAGOHB-207 | ENST00000544176.1 | 559 | - | IR | - | 162 | 59 | 338 | - | - | - | - |
| MAGOHB-208 | ENST00000544850.5 | 1057 | 90 | NMD | 62 | 62 + 94 | 59 | 111 | 9 + 277 | 83 | 362 | 722 |
| MAGOHB-209 | ENST00000545236.5 | 868 | 81 | NMD | 77 | 77 + 94 | 152 + 187 | 59 | 111 | 83 | 105 | 545 |
| MAGOHB-210 | ENST00000546173.5 | 576 | 81 | NMD | 41 | 41 + 94 | 152 + 12 | 59 | 111 | 107 | - | 289 |
| MAGOHB-211 | ENST00000625272.1 | 319 | 81 | Protein-coding | 73 | 73 + 94 | 152 | - | - | - | - | - |

\*NMD = Nonsense-mediated mRNA decay, IR= Intron retention, \* Number of bases written in red correspond to untranslated regions (UTR)

The *MAGOHB* gene in humans is located on the reverse strand of chromosome 12 at 12p13.2 *(chr12:10,604,193-10,613,609, GRCh38/hg38).* As per the Ensembl genome browser, *MAGOHB* pre-mRNA generates eleven alternately spliced transcripts (Table-1). Of these eleven transcripts, three have coding potential. Thus, similar to *MAGOH, MAGOHB* also produces multiple protein coding transcripts. Two of the protein-coding transcripts, *MAGOHB-201* and *MAGOHB-204*, contain five exons. *MAGOHB-201* is the primary transcript and codes for the 148 aa primary protein isoform, while the alternate transcript, *MAGOHB-204* codes for the alternate 102 aa protein isoform. The two transcripts differ due to the presence of an alternative 5’ splice site in the first exon of *MAGOHB-204* (Figure-1a). The third protein-coding transcript, *MAGOHB-211* consists of 2 exons which code for an 81 aa protein. In the transcript, *MAGOHB-211*, its first exon is similar to the principal isoform, *MAGOHB-201*, whereas its second exon differs completely due to the presence of an alternate 3’ splice site. *MAGOHB* thus produces three different protein isoforms. It will be fascinating to analyse whether the alternate protein isoforms are also generated in other species. Apart from these three protein coding transcripts, five other transcripts (*MAGOHB-205, −206, −208, −209, −210*) are also generated. Out of these five, three transcripts *MAGOHB-208, MAGOHB-209* and *MAGOHB-205* are made of six exons whereas *MAGOHB-206* and *MAGOHB-210* are composed of five exons. All the five transcripts are degraded via the NMD pathway due to the presence of premature termination codons (PTCs), which is one of the prerequisites of NMD. Lastly, the remaining three transcripts of *MAGOHB, MAGOHB-202, −203, −207* have intron retention (IR), and do not code for any protein. Here, *MAGOHB-203* and *MAGOHB-207* comprise of three exons whereas *MAGOHB-202* is made of two exons. In this article, we will focus only on the protein coding transcripts of both the MAGOH-paralogs.

In line with the Ensembl annotation, we will be referring to the protein isoforms according to their transcript IDs. We observed that the two MAGOH protein isoforms, principal isoform-146 aa and alternate isoform-109 aa differ in 37 residues, i.e., amino acids 50-86 of the principal isoform. In case of MAGOHB, three protein isoforms are generated. Coding region of the transcript *MAGOHB-204* starts from the second exon, and translates into an alternate protein isoform that varies from the principal protein isoform, MAGOHB-201 in the N-terminus (devoid of residues 1-46). The third protein isoform, MAGOHB-211 differs from the principal isoform in amino acids 38-81 since its second exon is formed by an alternate 3’ splice site. The presence of two functionally redundant MAGOH paralogs is in itself fascinating. From this characterisation we also know that both the MAGOH paralogs generate multiple protein isoforms in humans. In the following sections we have analysed the presence of multiple protein isoforms of MAGOH paralogs across different species, which might hint on an evolutionarily conserved function of the alternate protein isoforms generated by the MAGOH paralogs.

### 2.2 *MAGOHB* codes for multiple protein isoforms in mice

The MAGOH paralogs are conserved across vertebrates and MAGOH has been widely studied in mice. The presence of multiple protein isoforms of the MAGOH paralogs in humans raised the next question - are such alternate protein isoforms also present in mice? Hence, we analysed the mouse genome on Ensembl for multiple isoforms of MAGOH paralogs (Table-2). Here, we have referred to the mouse orthologs of human *MAGOH* paralogs as *Magoh* and *Magohb*. In mice, *Magoh* is located on chromosome 4 (*chr 4:107,879,755-107,887,424, GRCm38*), and is transcribed into two transcripts, *Magoh-201* and *Magoh-202. Magoh-201* is translated into a 146 aa protein, whereas *Magoh-202* does not code for any protein because of the absence of an ORF. Therefore, unlike humans, mice do not produce any alternate protein isoform of Magoh. The paralog of *Magoh, Magohb* is located on the reverse strand of chromosome 6 (*chr 6:131,284,388-131,293,244, GRCm38*). A total of seven *Magohb* transcripts are generated in mice, where two transcripts (*Magohb-201* and *Magohb-205*) are protein-coding. *Magohb-201* translates into the 148 aa principal protein isoform, whereas *Magohb-205* codes for the alternate protein isoform of 102 aa. Thus, in humans and mice two protein isoforms of MAGOHB are generated, which are of the same length. Similar to the alternate human MAGOHB transcript, *MAGOHB-204*, the first exon of *Magohb-205* constitutes most of the 5’UTR region and ligates exon 2 via an alternate 5’ splice site (Figure-1b). The start codon for *Magohb-205* is present in the second exon, thus the protein it encodes is devoid of residues 1 - 46 compared to the principal isoform. Curiously, splicing patterns of the two Magohb protein isoforms in mice are very similar to human MAGOHB protein isoforms. Besides the protein-coding transcripts, three transcripts, *Magohb-202, −203, −204,* are subjected to degradation via NMD. The remaining two transcripts (*Magohb-206* and *Magohb-207*) retain intronic regions and do not code for any protein. Thus, similar to its human ortholog, *Magohb* in mice also generates multiple protein-coding and non-coding splice variants.

**Table 2-.** Magoh paralog isoforms in *Mus musculus*

| Transcript ID | Ensembl ID | Length<br>(bases) | Protein<br>(amino acids) | Biotype | 5' UTR<br>(bases) | Exon 1<br>(bases) | Exon 2<br>(bases) | Exon3<br>(bases) | Exon4<br>(bases) | Exon5<br>(bases) | Exon6<br>(bases) | 3' UTR<br>(bases) |
| --- | --- | --- | --- | --- | --- | --- | --- | --- | --- | --- | --- | --- |
| **Magoh-201** | ENSMUST00000030348.5 | 692 | 146 | Protein coding | 103 | 103* + 88 | 59 | 111 | 83 | 100 + 148 | - | 148 |
| Magoh-202 | ENSMUST00000141376.1 | 483 | - | Processed transcript | - | 189 | 83 | 211 | - | - | - | - |
| **Magohb-201** | ENSMUST00000032307.11 | 767 | 148 | Protein coding | 79 | 79 + 94 | 59 | 111 | 83 | 100 + 241 | - | 241 |
| Magohb-202 | ENSMUST00000172883.7 | 460 | 35 | NMD* | - | 60 | 47 + 64 | 83 | 206 | - | - | 353 |
| Magohb-203 | ENSMUST00000173198.7 | 561 | 44 | NMD | - | 80 | 53 + 147 | 59 | 111 | 83 | 28 | 428 |
| **Magohb-204** | ENSMUST00000173332.1 | 395 | 44 | NMD | - | 80 | 54 + 147 | 59 | 56 | - | - | 262 |
| Magohb-205 | ENSMUST00000173837.5 | 897 | 102 | Protein coding | 564 | 520 | 44 + 15 | 111 | 83 | 100 + 24 | - | 24 |
| Magohb-206 | ENSMUST00000174488.1 | 447 | - | IR* | - | 231 | 216 | - | - | - | - | - |
| Magohb-207 | ENSMUST00000174781.3 | 546 | - | IR | - | 288 | 111 | 83 | 64 | - | - | - |

\*NMD = Nonsense-mediated mRNA decay, IR= Intron retention, \* Number of bases written in red correspond to untranslated regions (UTR)

Altogether, we find *MAGOH* transcript variants in humans and mice belong to two biotypes-protein coding transcripts and transcripts without an ORF; while *MAGOHB* transcript variants belong to three biotypes - protein-coding, NMD-sensitive, and IR. In both the species, *MAGOH* and *MAGOHB* generate multiple splice variants. The pattern of splicing for MAGOH-paralogs is similar, as both the species produce protein-coding as well as non-coding transcripts. In general, multiple splice variants tend to increase the proteome diversity of the genome. Among various alternative splicing (AS) events, IR, and AS-NMD are important for post-transcriptional gene regulation ^16,17^. It is also likely that transcripts with similar exon-intron structure, referred to as iso-orthologs, have similar biological function^18^. The similarity in splicing pattern as well as length of the alternate protein-coding transcripts prompted us to shift our focus to the sequence conservation of the alternate protein isoforms, discussed in the next section. The reason behind generation of multiple protein-coding and non-coding splice variants of MAGOH-paralogs is not yet known. They may contribute either towards regulation of MAGOH-paralogs or may perform some unexplored function.

### 2.3 Alternate isoforms of MAGOH paralogs are conserved across different species

We have described that MAGOH-paralogs produce more than one protein isoform (Figure-1). In this section, we depict the conserved nature of the alternate protein isoforms. To gain further insight into the conservation of the alternate proteins of MAGOH paralogs, we performed pairwise alignments of the protein sequences in humans and mice. We first compared the principal isoforms of MAGOH-paralogs, MAGOH-202 with Magoh-201 and MAGOHB-201 with Magohb-201, followed by comparison of the alternate isoforms of MAGOHB, MAGOHB-204 with Magohb-205.

The principal isoforms of MAGOH in humans and mice are 100 % identical (Figure-2a), whereas in the case of MAGOHB the identity is 98%, owing to three different residues, i.e., amino acids 2-4 (Figure-2b). Conservation of the primary isoforms is evident due to their suggested roles in EJC and NMD ^11,19^. Proceeding to the alternate protein isoforms, as discussed above, the alternate protein of MAGOH (MAGOH-201, 109 aa) does not have any ortholog in mice, meaning this isoform is either not important or might be specifically generated in higher vertebrates including humans. Interestingly, the 102 aa MAGOHB proteins in humans and mice are 100% identical to each other (Figure-2c). The striking identity prompted us to analyze other species for the presence of alternate isoforms of MAGOH-paralogs. Thus, we analyzed the alternate isoforms in different species and performed a multiple sequence alignment of the protein sequence. Quite interestingly, the alternate isoforms of the MAGOH-paralogs are conserved across vertebrates (Table 3). Though the alternate isoform of MAGOH, 109 aa is not present in mice, it is present in 52 other species, where the protein sequence differs only at position 2 (Supplementary Figure-1 and 2). The alternate isoform of MAGOHB, 102 aa, is present in 11 different species and is 100% identical in 10 species, but differs in the alternate isoform of kangaroo rat-*Dipodomys ordii* at position 25 and 35. We have also validated the expression of the conserved alternate protein-coding transcripts, *MAGOH-201* and *MAGOHB-204* in HEK-293 cells via RT-PCR (Figure-3a). We further performed a phylogenetic analysis of cDNA sequence from four species having alternate isoforms for both MAGOH and MAGOHB (highlighted in Table 3). The tree estimated via maximum likelihood shows the isoforms clustered into two groups, specific for MAGOH and MAGOHB. This indicates that similar to the principal isoforms, the alternate isoforms are also evolutionarily conserved orthologs (Figure-3b). The conservation of the alternate protein isoforms of MAGOH paralogs across species is quite riveting. Consequently, an evolutionary pressure must have resulted in conservation of the alternate protein isoforms of both the MAGOH-paralogs. It has been previously pointed that an identity in the range of 50%-70% is the defining boundary of protein conservation, and function of conserved proteins is generally expected to be preserved above 70% identity ^20^. Thus, the presence of a conserved alternate isoform of MAGOH and MAGOHB does not seem to be a mere coincidence and might involve a functional role, yet unexplored. Due to the difference in the residues of the principal and alternate isoforms of MAGOH paralogs, there might exist structural and functional differences between them which we have elaborated below.

**Figure 2-.**
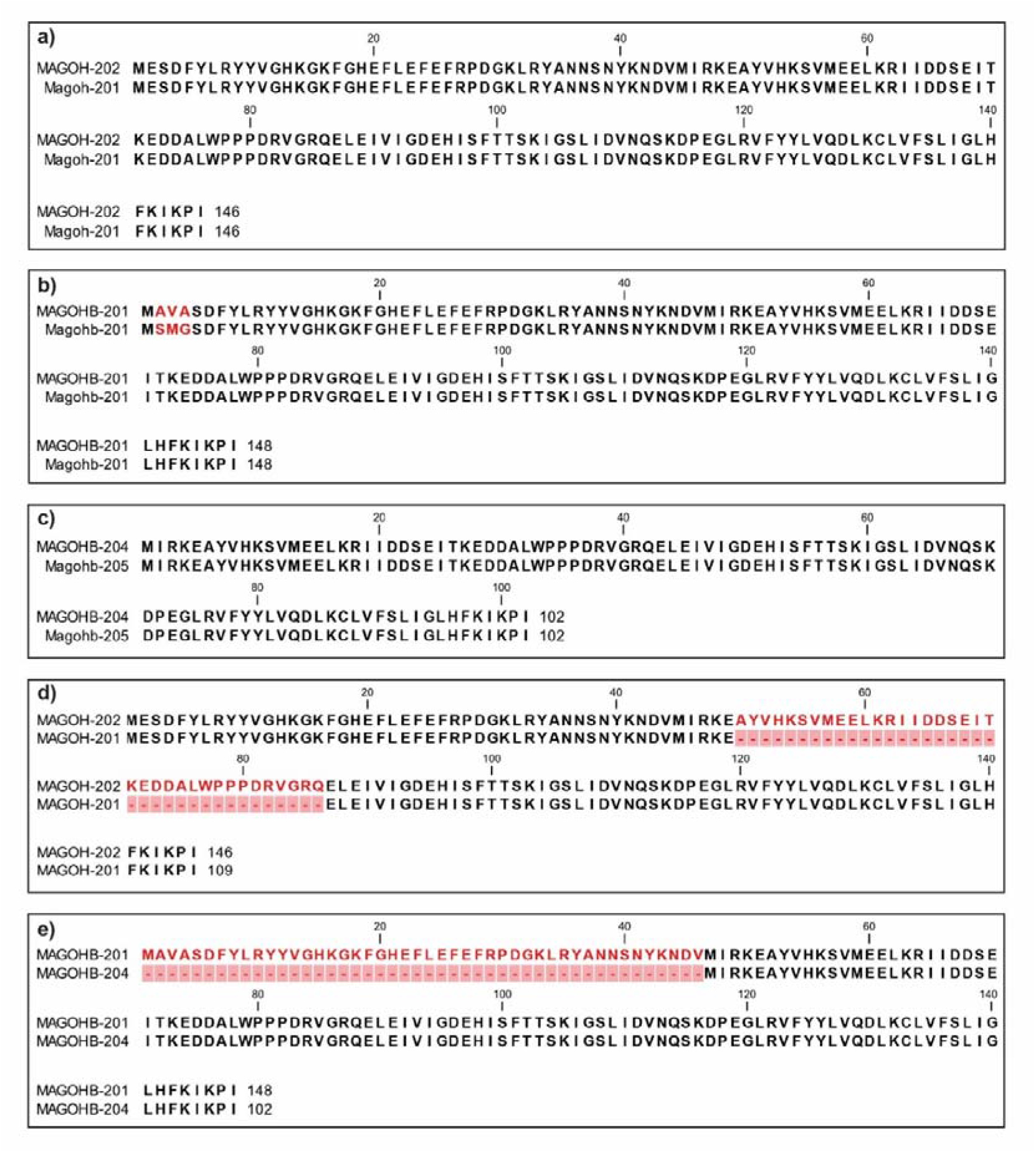
Pairwise alignment of MAGOH protein isoforms. The protein isoform alignments were generated using the CLC Main Workbench. **a)** MAGOH principal isoform alignment in humans (MAGOH-202) and mice (Magoh-201) **b)** MAGOHB principal isoform alignment in humans (MAGOHB-201) and mice (Magohb-201) **c)** MAGOHB alternate isoform alignment in humans (MAGOHB-204) and mice (Magohb-205). The principal and alternate isoforms of MAGOH/MAGOHB in humans were also aligned to each other to analyse similarity - **d)** MAGOH principal isoform (MAGOH-202) aligned with alternate isoform (MAGOH-201) **e)** MAGOHB principal isoform (MAGOHB-201) aligned with its alternate isoform (MAGOHB-204). The different residues are coloured in red, gaps are coloured in pink boxes and represented as dashes.

**Figure 3-.**
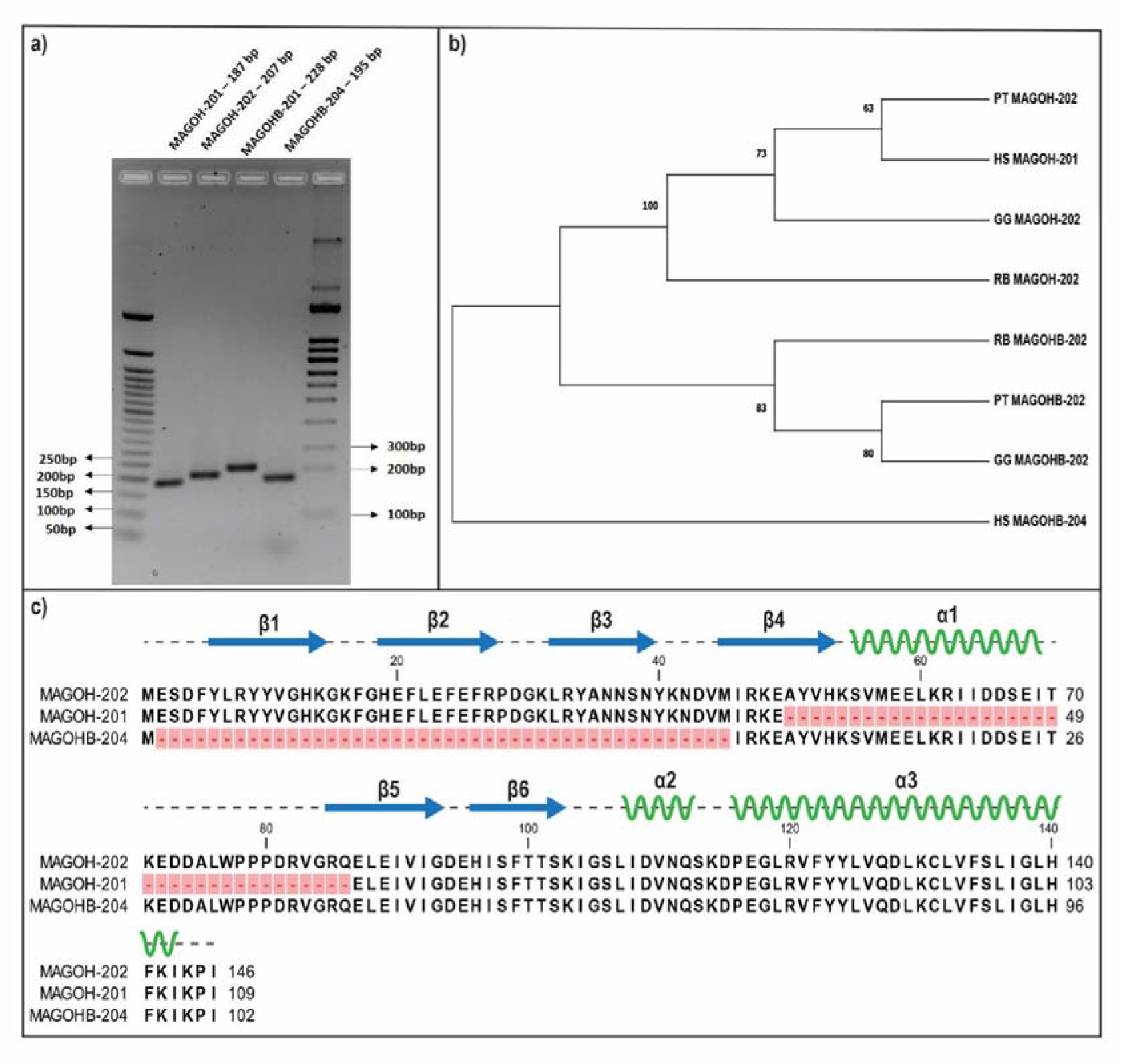
Analysis of alternate isoforms of MAGOH paralogs. **a)** The expression of alternate isoforms was validated in HEK-293 cells. The expected size of amplicons is mentioned in brackets. **b)** Phylogenetic tree for species expressing both the isoforms of MAGOH paralogs. The numbers on each node indicate bootstrap values for 500 replicates. PT-*Pan troglodytes*, HS-*Homo sapiens*, GG-*Gorilla gorilla*, RB-*Rhinopithecus bieti*. **c)** Structural comparison of alternate isoforms to the principal MAGOH isoform, MAGOH-202. MAGOH-202 is aligned with MAGOH-201 (109 amino acids, devoid of residues 50-86) and MAGOHB-204 (102 amino acids, lacking N-terminus residues 1-46). The gaps are represented in pink boxes. Secondary structures are represented above the alignment according to the crystal structure of MAGOH-Y14 (PDB ID 1P27), described by Lau et al^15^. Here, β1-6 represent β strands and α1-3 represent α-helices.

**Table 3-.** Species with conserved alternate isoforms of MAGOH paralogs

| Gene | Size of protein<br>(amino acids) | Species with conserved isoform |
| --- | --- | --- |
| MAGOH | 109 | *Anser brachyrhynchus*, *Anser cygnoides*, *Aotus nancymae*, *Bison bison bison*, *Bos grunniens*, Hybrid - *Bos Indicus (Bos indicus x Bos taurus)*, *Bos mutus*, Hybrid - *Bos Taurus (Bos indicus x Bos taurus)*, *Callithrix jacchus*, *Camelus dromedarius*, *Canis lupus familiaris*, *Canis lupus familiaris*, *Carlito syricta*, *Catagonus wagneri*, *Cebus capucinus imitator*, *Cercocebus atys*, *Colobus angolensis palliatus*, *Equus caballus*, *Felis catus*, ***Gorilla gorilla gorilla***, ***Homo sapiens***, *Lynx canadensis*, *Macaca fascicularis*, *Macaca nemestrina*, *Mandrillus leucophaeus*, *Microcebus murinus*, *Monodelphis domestica*, *Moschus moschiferus*, *Neovison vison*, *Nomascus leucogenys*, *Ornithorhynchus anatinus*, *Oryctolagus cuniculus*, *Bonobo*, *Pan paniscus*, ***Pan troglodytes***, *Panthera leo*, *Panthera pardus*, *Panthera tigris altaica*, *Papio anubis*, *Physeter catodon*, *Piliocolobus tephrosceles*, *Podarcis muralis*, *Prolemur simus*, *Propithecus coquereli*, *Rhinolophus ferrumequinum*, ***Rhinopithecus bieti***, *Rhinopithecus roxellana*, *Saimiri boliviensis boliviensis*, *Suricata suricatta*, *Sus scrofa*, *Theropithecus gelada*, *Ursus thibetanus thibetanus*, *Vombatus ursinus* |
| MAGOHB | 102 | *Cavia porcellus, Cricetulus griseus, Dipodomys ordii, **Gorilla gorilla gorilla, Homo sapiens, Mus musculus, Mus pahari, Mus spretus, Nannospalax galili, Pan troglodytes, Rhinopithecus bieti*** |

The structure of MAGOH is composed of three α helices and six β strands that form an extended sheet^15^. The helices form a hydrophobic core to interact with the RNA-binding domain of Y-14, which is primarily involved in protein-protein interactions. The extended β sheet contains conserved residues, forming the binding platform for EJC factors or other associated proteins ^15,19^. On aligning the principal and alternate protein isoforms in humans, we found the alternate isoform of MAGOH to have 74.7% sequence identity to the principal MAGOH isoform, whereas the alternate MAGOHB isoform has 68.9% sequence identity to the principal MAGOHB isoform (Figure-2d, 2e). Comparison of structural features is shown in Figure-4. As shown, the alternate isoform of MAGOH lacks residues 50-86 corresponding to the principal MAGOH isoform, MAGOH-202. These residues constitute a portion of the α_1_ helix of MAGOH. Helix α_1_ with helix α_3_ form a part of the hydrophobic core which binds the RNA-binding domain (RBD) of Y-14. Consequently, the alternate isoform, MAGOH-201, might lack the ability to bind Y-14 or may bind to it without masking Y-14’s RBD. If the alternate isoform of MAGOH binds Y-14 it will be interesting to find out the stability and possible function of this alternate heterodimer complex.

The N-terminus of the principal MAGOH isoform contains conserved residues, Tyr6, Tyr10, and Phe21 of the β-sheet^15^. Since the alternate isoform of MAGOHB, MAGOHB-204, lacks residues 1-46, it also lacks these conserved residues present in the β sheet. Consequently, mutations in residues 16/17, 20, 39/40, 41/42 of the principal MAGOH protein result in loss of interaction with pre-mRNA, spliced mRNA as well as EJC components, BTZ, UPF3b and eIF4A3, whereas interaction with Y-14 and PYM remains intact^21^. Thus, MAGOHB-204 might have functional Y-14 binding but may lack binding with other EJC factors.

To summarize, we find that the generation of alternate protein isoforms by MAGOH-paralogs is not only limited to humans, but also extends to other species. The high degree of conservation of the alternate protein isoforms, MAGOH-201 (109 aa) and MAGOHB-204 (102 aa), may point out to the presence of an evolutionary pressure maintaining multiple isoforms of MAGOH and MAGOHB. These alternate protein isoforms might be involved in regulating the levels of principal MAGOH proteins or may have functions entirely different from the principal isoforms.

Delineating the function of these isoforms is challenging because of the sequence similarity between the primary and alternate protein isoforms. Hence, the function of the alternate isoforms might need to be analysed in conditions where the principal isoform is absent, and is the subject of further experimentations.

## 3. MATERIALS AND METHODS

### 3.1 Sequence information for isoforms

All the sequence information for *Homo sapiens* and *Mus musculus* were taken from Ensembl (Release 101) (https://asia.ensembl.org/index.html) ^22^. The Ensembl IDs are - ENSG00000162385(MAGOH), ENSG00000111196(MAGOHB) for human MAGOH paralogs, and ENSMUSG00000028609(Magoh), ENSMUSG00000030188(Magohb) for Magoh paralogs in mice. Genome assembly versions referred were Human-GRCh38.p13 and Mouse-GRCm38.p6.

### 3.2 Sequence alignment

Alignments were generated using the CLC Main Workbench’s alignment tool and the identity among aligned proteins was analysed using the EMBOSS Needle pairwise alignment tool^23^. The proteins were aligned with the BLOSUM62 matrix. Gap opening penalty was kept at 10 and gap extension penalty at 0.5. Multiple sequence alignments of the alternate isoforms (109 aa and 102 aa) were performed using ClustalW in the MEGA-X^24^ tool with the following parameters-gap opening penalty 10.00, gap extension penalty 0.20. Jalview was used for visualisation of the multiple sequence alignments^25^. Phylogenetic trees were generated using MEGA-X as previously described^26^, briefly the cDNA sequence was downloaded from Ensembl, codons aligned using MUSCLE followed by model generation and using the highest model to estimate the phylogenetic tree via maximum likelihood.

### 3.3 Isoform expression analysis

RNA for isoform analysis was extracted from HEK-293 cells, cultured in DMEM at 37°C in a humidified CO_2_ incubator. The RNA was converted into cDNA using the Super Reverse Transcriptase MuLV cDNA kit (Biobharati). The primers sequence for the principal isoform were used as described previously^11^. Primer sequences for the alternate isoforms are mentioned in the supplementary file.

## ACKNOWLEDGEMENT

We would like to thank all the members of the KKS lab for their helpful suggestions, especially Bhagyashree Deka for her insightful comments on the manuscript. A.R. and P.C. are funded under the scholarships provided by Ministry of Human Resource Development (MHRD), Govt. of India. K.K.S. acknowledges funding from the Science & Engineering Research Board (SERB), Government of India [Grant No. CRG/2019/001352]. We would also like to thank the Indian Institute of Technology, Guwahati for providing required resources.

## Abbreviations

EJC: Exon Junction Complex
MAGOH: mago nashi homolog
eIF4A3: eukaryotic initiation factor 4 A3
RBM8A: RNA binding motif 8A (also known as Y-14)

## REFERENCES

1. Singh G, Kucukural A, Cenik C, Leszyk JD, Shaffer SA, Weng Z, Moore MJ. The cellular EJC interactome reveals higher-order mRNP structure and an EJC-SR protein nexus. Cell 151, 750–764 (2012). doi: 10.1016/j.cell.2012.10.007

2. Tange TØ, Shibuya T, Jurica MS, Moore MJ. Biochemical analysis of the EJC reveals two new factors and a stable tetrameric protein core. RNA 11, 1869–1883 (2005). doi: 10.1261/rna.2155905.

3. Kataoka N, Diem MD, Kim VN, Yong J, Dreyfuss G. Magoh, a human homolog of Drosophila mago nashi protein, is a component of the splicing-dependent exon-exon junction complex. EMBO J. 20, 6424–33 (2001). doi: 10.1093/emboj/20.22.6424.

4. Wang Z, Ballut L, Barbosa I, Le Hir H. Exon Junction Complexes can have distinct functional flavours to regulate specific splicing events. Sci Rep. 8, 9509 (2018). doi: 10.1038/s41598-018-27826-y.

5. Le Hir H, Izaurralde E, Maquat LE, Moore MJ. The spliceosome deposits multiple proteins 20–24 nucleotides upstream of mRNA exon-exon junctions. EMBO J. 19, 6860–9 (2000). doi: 10.1093/emboj/19.24.6860.

6. Le Hir H, Gatfield D, Braun IC, Forler D, Izaurralde E. The protein Mago provides a link between splicing and mRNA localization. EMBO Rep. 2, 1119–24 (2001). doi: 10.1093/embo-reports/kve245.

7. McMahon JJ, Miller EE, Silver DL. The exon junction complex in neural development and neurodevelopmental disease. Int J Dev Neurosci. 55, 117–123 (2016). doi: 10.1016/j.ijdevneu.2016.03.006. Epub 2016 Apr 9. PMID: 27071691.

8. Boswell RE, Prout ME, Steichen JC. Mutations in a newly identified Drosophila melanogaster gene, mago nashi, disrupt germ cell formation and result in the formation of mirror-image symmetrical double abdomen embryos. Development. 113, 373–84 (1991). Erratum in: Development 1991 Dec;113(4):precedi. PMID: 1765008.

9. Gehring NH, Kunz JB, Neu-Yilik G, Breit S, Viegas MH, Hentze MW, Kulozik AE. Exon-junction complex components specify distinct routes of nonsense-mediated mRNA decay with differential cofactor requirements. Molecular cell. 20, 65–75 (2005). doi: 10.1016/j.molcel.2005.08.012. PMID: 16209946.

10. Gangras P, Gallagher TL, Parthun MA, Yi Z, Patton RD, Tietz KT, Deans NC, Bundschuh R, Amacher SL, Singh G. Zebrafish rbm8a and magoh mutants reveal EJC developmental functions and new 3’UTR intron-containing NMD targets. PLoS genetics. 16, e1008830 (2020). doi: 10.1371/journal.pgen.1008830. PMID: 32502192; PMCID: PMC7310861.

11. Singh KK, Wachsmuth L, Kulozik AE, Gehring NH. Two mammalian MAGOH genes contribute to exon junction complex composition and nonsense-mediated decay. RNA Biology 10, 1291–1298 (2013).

12. Viswanathan SR, Nogueira MF, Buss CG, Krill-Burger JM, Wawer MJ, Malolepsza E, Berger AC, Choi PS, Shih J, Taylor AM, Tanenbaum B, Pedamallu CS, Cherniack AD, Tamayo P, Strathdee CA, Lage K, Carr SA, Schenone M, Bhatia SN, Vazquez F, Tsherniak A, Hahn WC, Meyerson M. Genome-scale analysis identifies paralog lethality as a vulnerability of chromosome 1p loss in cancer. Nature Genetics 50, 937–943 (2018). doi: 10.1038/s41588-018-0155-3. PMID: 29955178; PMCID: PMC6143899.

13. Zhou Y, Li Z, Wu X, Tou L, Zheng J, Zhou D. MAGOH/MAGOHB Inhibits the Tumorigenesis of Gastric Cancer via Inactivation of b-RAF/MEK/ERK Signaling. OncoTargets and therapy 13, 12723–12735 (2020). doi: 10.2147/OTT.S263913. PMID: 33328743; PMCID: PMC7735944.

14. Zhao XF, Colaizzo-Anas T, Nowak NJ, Shows TB, Elliott RW, Aplan PD. The mammalian homologue of mago nashi encodes a serum-inducible protein. Genomics. 47, 319–22 (1998). doi: 10.1006/geno.1997.5126.

15. Lau CK, Diem MD, Dreyfuss G, Van Duyne GD. Structure of the Y14-Magoh core of the exon junction complex. Curr Biol. 13, 933–41 (2003). doi: 10.1016/s0960-9822(03)00328-2.

16. McGlincy NJ, Smith CW. Alternative splicing resulting in nonsense-mediated mRNA decay: what is the meaning of nonsense? Trends Biochem Sci. 33, 385–93 (2008). doi: 10.1016/j.tibs.2008.06.001.

17. Jacob AG, Smith CWJ. Intron retention as a component of regulated gene expression programs. Hum Genet. 136, 1043–1057 (2017). doi: 10.1007/s00439-017-1791-x.

18. Zambelli F, Pavesi G, Gissi C, Horner DS, Pesole G. Assessment of orthologous splicing isoforms in human and mouse orthologous genes. BMC Genomics. 11, 534 (2010). doi: 10.1186/1471-2164-11-534.

19. Fribourg S, Gatfield D, Izaurralde E, Conti E. A novel mode of RBD-protein recognition in the Y14-Mago complex. Nat Struct Biol. 10, 433–9 (2003). doi: 10.1038/nsb926.

20. Morata J, Béjar S, Talavera D, Riera C, Lois S, de Xaxars GM, de la Cruz X. The relationship between gene isoform multiplicity, number of exons and protein divergence. PLoS One. 8, e72742 (2013). doi: 10.1371/journal.pone.0072742.

21. Gehring NH, Lamprinaki S, Hentze MW, Kulozik AE. The hierarchy of exon-junction complex assembly by the spliceosome explains key features of mammalian nonsense-mediated mRNA decay. PLoS Biol. 7 e1000120 (2009). doi: 10.1371/journal.pbio.1000120.

22. Yates AD, Achuthan P, Akanni W, Allen J, Allen J, Alvarez-Jarreta J, Amode MR, Armean IM, Azov AG, Bennett R, Bhai J, Billis K, Boddu S, Marugán JC, Cummins C, Davidson C, Dodiya K, Fatima R, Gall A, Giron CG, Gil L, Grego T, Haggerty L, Haskell E, Hourlier T, Izuogu OG, Janacek SH, Juettemann T, Kay M, Lavidas I, Le T, Lemos D, Martinez JG, Maurel T, McDowall M, McMahon A, Mohanan S, Moore B, Nuhn M, Oheh DN, Parker A, Parton A, Patricio M, Sakthivel MP, Abdul Salam AI, Schmitt BM, Schuilenburg H, Sheppard D, Sycheva M, Szuba M, Taylor K, Thormann A, Threadgold G, Vullo A, Walts B, Winterbottom A, Zadissa A, Chakiachvili M, Flint B, Frankish A, Hunt SE, IIsley G, Kostadima M, Langridge N, Loveland JE, Martin FJ, Morales J, Mudge JM, Muffato M, Perry E, Ruffier M, Trevanion SJ, Cunningham F, Howe KL, Zerbino DR, Flicek P. Ensembl 2020. Nucleic Acids Res. 48, D682–D688 (2020). doi: 10.1093/nar/gkz966.

23. Madeira F, Park YM, Lee J, Buso N, Gur T, Madhusoodanan N, Basutkar P, Tivey ARN, Potter SC, Finn RD, Lopez R. The EMBL-EBI search and sequence analysis tools APIs in 2019. Nucleic Acids Res. 47, W636–W641 (2019). doi: 10.1093/nar/gkz268.

24. Kumar S, Stecher G, Li M, Knyaz C, Tamura K. MEGA X: Molecular Evolutionary Genetics Analysis across Computing Platforms. Mol Biol Evol. 35, 1547–1549 (2018). doi: 10.1093/molbev/msy096.

25. Waterhouse AM, Procter JB, Martin DMA, Clamp M, Barton GJ. Jalview Version 2 - a multiple sequence alignment editor and analysis workbench. Bioinformatics 25, 1189–1191 (2009).doi: 10.1093/bioinformatics/btp033s

26. Barry G Hall. Building Phylogenetic Trees from Molecular Data with MEGA. Molecular Biology and Evolution 30, 1229–1235 (2013). doi: 10.1093/molbev/mst012.

