## Supplementary Files for "The MAGOH paralogs - MAGOH, MAGOHB and their multiple isoforms"

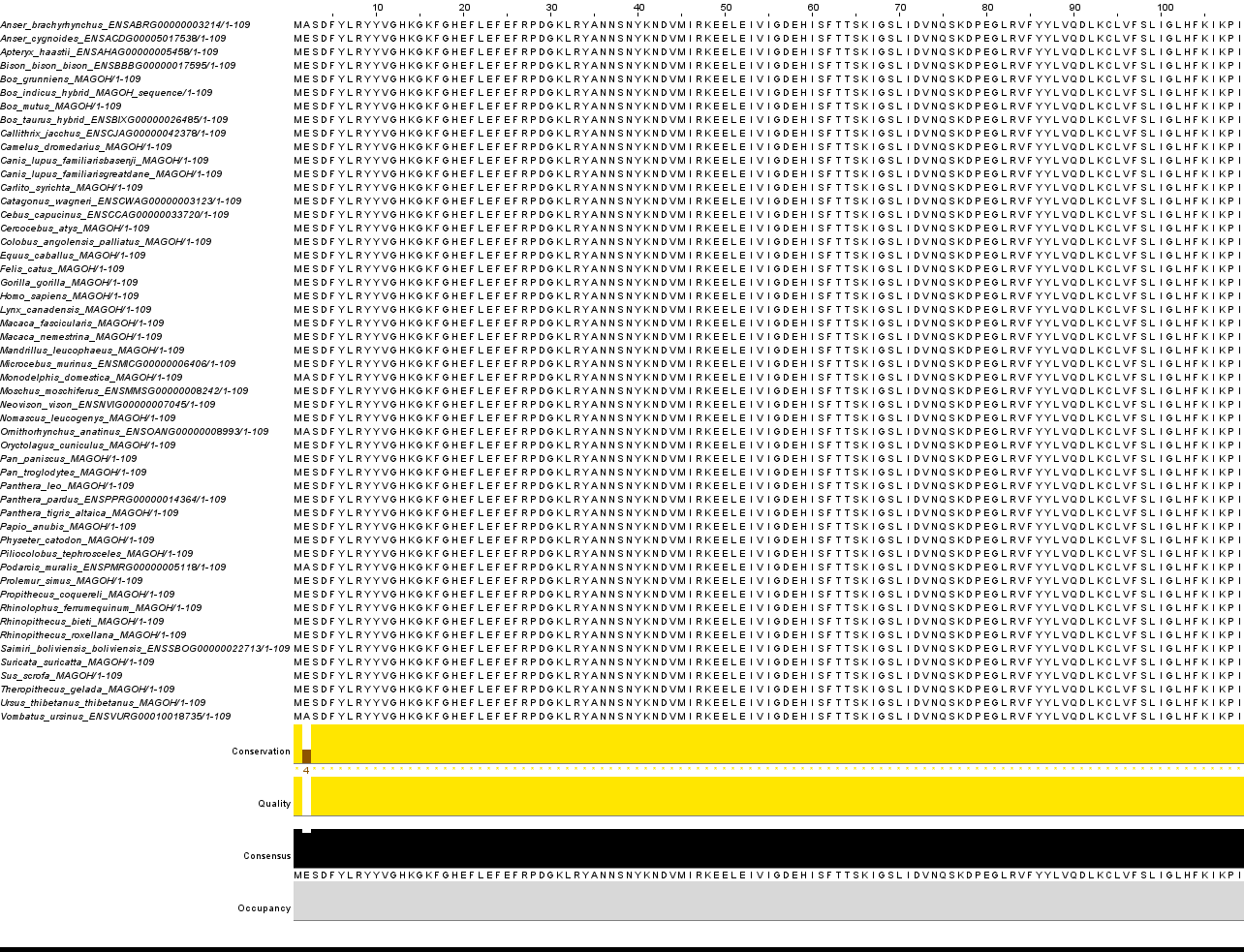

**Supplementary Figure-1**: Multiple Sequence alignment of the protein sequence of the alternate MAGOH isoform, MAGOH-201. The sequence across 52 species is shown, where only position 2 is not conserved.

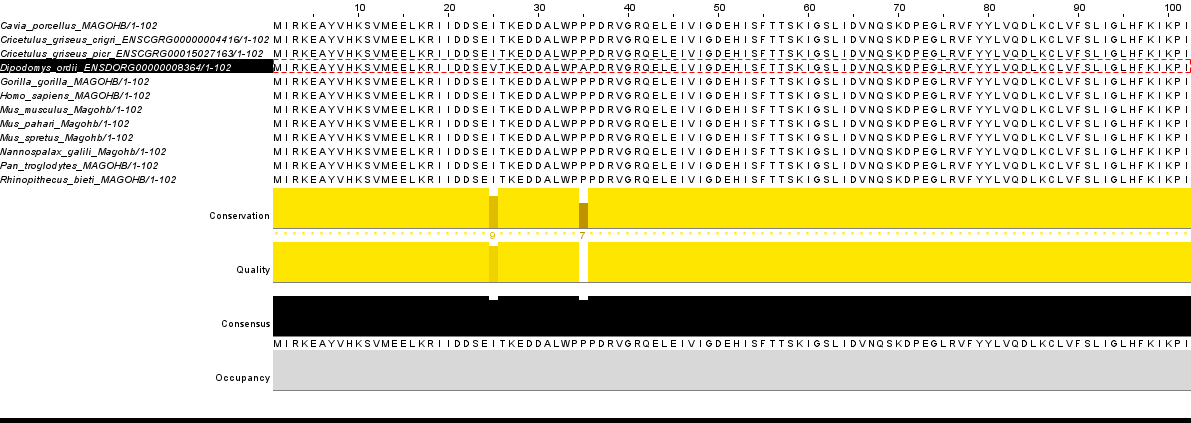

**Supplementary Figure-2**: Multiple Sequence alignment of the protein sequence of the conserved alternate MAGOHB isoform, MAGOHB-204. The alignment across 11 species is shown. The 102 aa protein is identical across 10 species, and differs in the sequence of *Dipodomys ordii* at position 25 and 35.

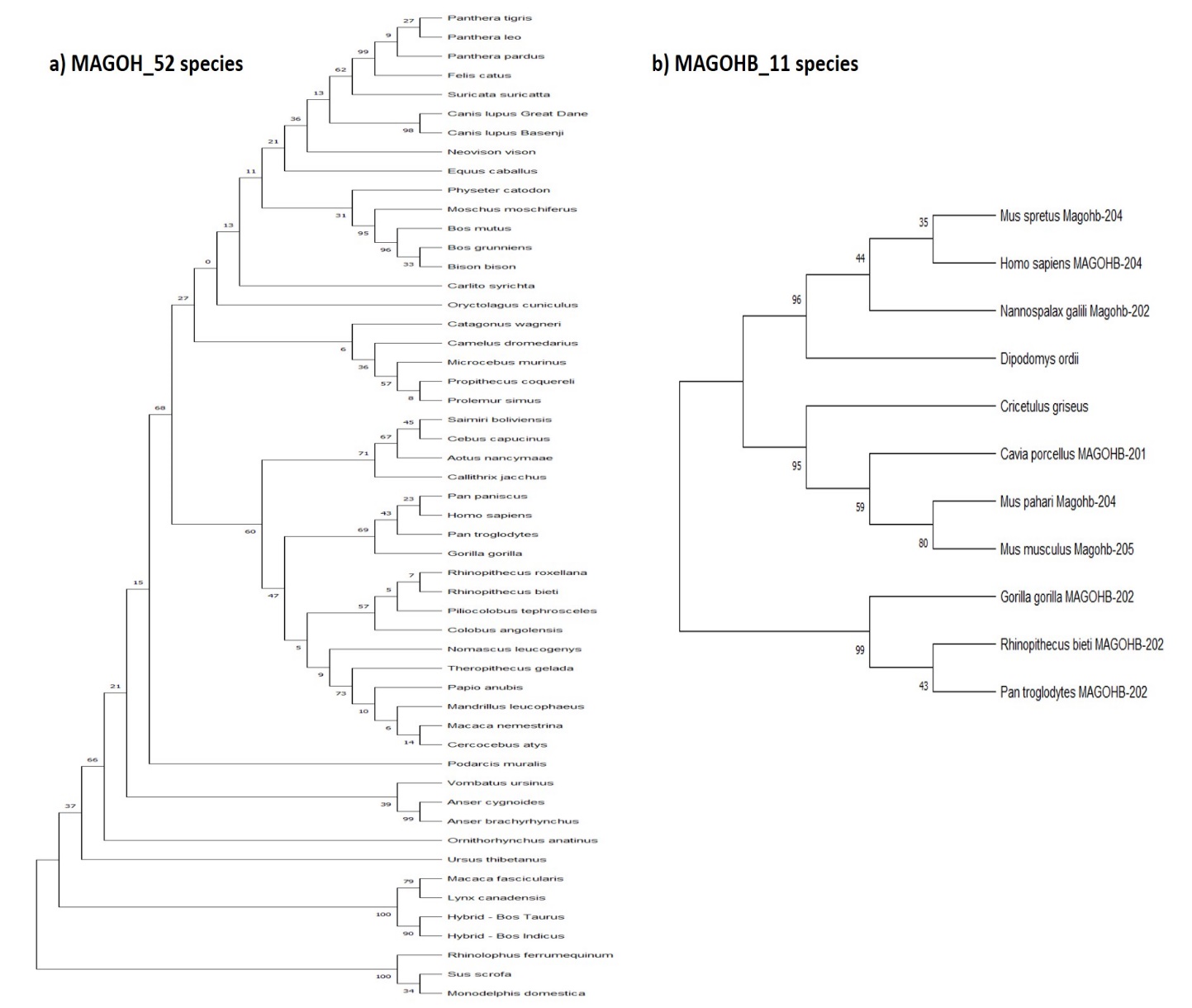

**Supplementary Figure-3**: Phylogenetic trees for species referred to in Table-3 of the main manuscript. The evolutionary histories were inferred by using the Maximum Likelihood method and Tamura 3-parameter model^1^. Evolutionary analyses were conducted in MEGA X^2^. Bootstrapping was performed for 500 replicates. The percentage of trees in which the associated taxa clustered together is shown next to the branches. **a)** For 109 aa alternate isoform of MAGOH in 52 species, tree with the highest log likelihood (-3385.37) is shown. **b)** For 102aa alternate isoform of MAGOHB in 11 species, the tree with the highest log likelihood (-2166.29) is shown

**Supplementary Table 1 –** Primer sequences used to amplify alternate isoforms

| **Isoform** | **Primer Sequences** |
| --- | --- |
| MAGOH-201 | Forward Primer- 5’-TGCCTTGCAGATCCTCCTG-3’ |
|  | Reverse Primer- 5’- CGATTTCAAGCTCCTCTTTTCTG -3’ |
| MAGOHB-204 | Forward Primer-5’-GCTTCCCGGGAAAGCTTAGAT-3’ |
|  | Reverse Primer- 5’-GCTTCCCGGGAAAGCTTAGAT-3’ |

**References**

1. Tamura K. (1992). Estimation of the number of nucleotide substitutions when there are strong transition-transversion and G + C-content biases. Molecular Biology and Evolution 9:678-687.

2. Kumar S., Stecher G., Li M., Knyaz C., and Tamura K. (2018). MEGA X: Molecular Evolutionary Genetics Analysis across computing platforms. Molecular Biology and Evolution 35:1547-1549.
